# Nor-LAAM loaded PLGA Microparticles for Treating Opioid Use Disorder

**DOI:** 10.1101/2024.04.08.588574

**Authors:** Diane Ingabire, Chaolong Qin, Tuo Meng, Aji Alex Moothendathu Raynold, Hadi Sudarjat, E. Andrew Townsend, Rudra Pangeni, Sagun Poudel, Michelle Arriaga, Long Zhao, Woon. N Chow, Matthew Banks, Qingguo Xu

## Abstract

The treatment landscape for opioid use disorder (OUD) faces challenges stemming from the limited efficacy of existing medications, poor adherence to prescribed regimens, and a heightened risk of fatal overdose post-treatment cessation. Therefore, there is a pressing need for innovative therapeutic strategies that enhance the effectiveness of interventions and the overall well-being of individuals with OUD. This study explored the therapeutic potential of nor-Levo-α-acetylmethadol (nor-LAAM) to treat OUD. We developed sustained release nor-LAAM-loaded poly (lactic-co-glycolic acid) (PLGA) microparticles (MP) using a hydrophobic ion pairing (HIP) approach. The nor-LAAM-MP prepared using HIP with pamoic acid had high drug loading and exhibited minimal initial burst release and sustained release. The nor-LAAM-MP was further optimized for desirable particle size, drug loading, and release kinetics. The lead nor-LAAM-MP (F4) had a relatively high drug loading (11 wt.%) and an average diameter (19 µm) and maintained a sustained drug release for 4 weeks. A single subcutaneous injection of nor-LAAM-MP (F4) provided detectable nor-LAAM levels in rabbit plasma for at least 15 days. We further evaluated the therapeutic efficacy of nor-LAAM-MP (F4) in a well-established fentanyl-addiction rat model, and revealed a marked reduction in fentanyl choice and withdrawal symptoms in fentanyl-dependent rats. These findings provide insights into further developing long-acting nor-LAAM-MP for treating OUD. It has the potential to offer a new effective medication to the existing sparse armamentarium of products available to treat OUD.

## 1. Introduction

The United States is currently grappling with an unrelenting opioid crisis, with a concerning rise in opioid overdose deaths. The crisis is primarily fueled by the widespread use of mu-opioid receptor (MOR) agonists, particularly fentanyl [1]. The increase in opioid prescriptions from 2000 to 2013, reaching almost 207 million, made the United States the leading global consumer [2]. This surge correlated with a rise in opioid-related deaths, emphasizing the necessity for controlled prescribing practices. However, restrictions on prescribing opioids led to increased abuse of heroin and illicit fentanyl [3,4], which contributed to opioid overdose deaths. In 2021, 100,306 overdose deaths were recorded [5], and this underscores the critical need to implement effective intervention strategies.

Currently, methadone, buprenorphine, and naltrexone stand as the three FDA-approved medications for opioid use disorder (OUD) due to their proven efficacy. Methadone, a full MOR agonist, has demonstrated efficacy in reducing illicit opioid use and overdose deaths among OUD patients [6]. Buprenorphine and naltrexone, acting as partial agonists and antagonists to MOR, respectively, have played roles in reducing opioid craving, mitigating withdrawal symptoms, and preventing relapse [6–8]. Despite the effectiveness of Medications for OUD (MOUDs) [9,10], challenges persist, including poor patient retention, increased overdose risks, low efficacy [6,11], and undesired side effects [12,13]. Due to lower patient compliance with current treatments, sustained-release formulations of buprenorphine and naltrexone have been developed, which addressed some of these challenges [7,8]. Efforts to expand OUD medication availability include the use of methadone vans or mobile medication units and making treatment more accessible [14]. Nevertheless, cases of relapse and a population of patients unresponsive to the current OUD treatments remain a challenge [15].

Given these challenges in OUD treatment and the escalating crisis, urgent preclinical research is needed to develop more effective pharmacotherapies. The present study explored the therapeutic potential of nor-Levo-α-acetylmethadol (nor-LAAM), an active metabolite of Levo-α-acetylmethadol (LAAM) with even higher potency and better safety than LAAM. The published receptor binding studies showed that nor-LAAM has a higher binding affinity and is more potent at mu-opioid receptors compared to LAAM, with a binding affinity (K_i_) reported as 5.6 nM and 740 nM and potency (IC_50_) of 1.2 nM and 100,000 nM, for nor-LAAM and LAAM, respectively [16,17]. Furthermore, nor-LAAM (IC_50_=12 µM) is much less potent than LAAM (IC_50_=3 µM) and similar potency to methadone (IC_50_=10 µM) to inhibit human ether-a-go-go-related gene (hERG) channels associated with cardiac toxicity, indicating that nor-LAAM may represent an improved OUD treatment alternative to methadone and LAAM because of the wider therapeutic window [18]. Prior studies have shown that drug-loaded PLGA microparticles provide sustained drug release, easy parenteral administration, and are biocompatible/safe [19,20]. In the field of substance use disorder, PLGA polymers have been used and approved as marketed formulations like Vivitrol^®^ and Sublocade^®^ for alcohol and opioid dependence, respectively. In this study, we sought to explore the therapeutic potential of nor-LAAM for OUD by developing a sustained release polymeric microparticle (MP) formulation suitable for subcutaneous (SQ) injection, potentially beneficial for OUD patients who are unresponsive to methadone or buprenorphine treatments [21].

Nor-LAAM was available as HCl salt (nor-LAAM.HCl) with high water-solubility, and we found that it was very challenging to effectively load nor-LAAM.HCl into PLGA microparticles leading to low drug loading and short drug release duration. Therefore, we applied the hydrophobic ion pairing (HIP) method to improve drug loading and release [22]. The novel nor-LAAM MP through the HIP method exhibited promising attributes of high drug loading and sustained drug release suitable for parenteral administration. We tested the efficacy of lead nor-LAAM MP following a single SQ injection in fentanyl-dependent rats using a self-drug administration method, which is a widely studied preclinical approach to induce fentanyl dependence and evaluate withdrawal symptoms [23,24]. We believe that a sustained-release nor-LAAM-MP formulation is advantageous as it can enhance drug bioavailability, minimize adverse effects, reduce dosing frequency, and improve patient adherence [25].

## 2. Materials and methods

### 2.1. Materials

Poly (D, L-lactic-co-glycolic acid, LA: GA 50:50, Mw 5, 18, 34, 54 kDa acid terminated, Mw 54 kDa ester terminated) (PLGA) were obtained from Evonik Industries (Birmingham, AL). Nor-Levo-α-acetylmethadol Hydrochloride (nor-LAAM.HCl) and Fentanyl-HCl were sourced from the National Institute of Drug Abuse (NIDA) Drug Supply Program (Bethesda, MD). Ketoprofen 100 mg/ml was obtained from Bimeda USA (Oakbrook Terrace, IL). We obtained a 100 mg/ml Gentamicin sulfate solution from Aspen Veterinary Resources LTD (Loveland, CO). Heparin Sodium Injection USP units/mL was obtained from McKesson Corporation. 0.9% Sodium Chloride 50 mL from ICU Medical, Inc (Lake Forest, IL). Poly (vinyl alcohol) (PVA) with an average MW of 30,000-70,000, 87-90% hydrolysis level, and sodium docusate were obtained from Sigma-Aldrich (St. Louis, MO). Rabbit plasma was purchased from Biochemed Services (Winchester, VA). Oleic acid, EDTA (0.5M), 40-μm cell strainer, and 0.22 μm hydrophilic/hydrophobic filter were purchased from Thermo Fisher Scientific (Hanover Park, IL). Pamoic acid disodium monohydrate was obtained from TCI (Portland, OR). All other chemical reagents and solvents were analytical grade or above.

### 2.3. Preparation of nor-LAAM loaded PLGA microparticles

Nor-LAAM-loaded PLGA microparticles (Nor-LAAM-MP) without HIP were prepared using an emulsion solvent evaporation method [26,27]. In short, nor-LAAM.HCl and PLGA polymers (targeted drug loading 20 - 40 wt%) were dissolved in DCM. The mixture was added into 40 mL of a 1% (w/v) aqueous PVA solution under homogenization at 7000 rpm for 2 minutes using a High Shear mixer (Silverson, East Longmeadow, MA). The emulsion was transferred to a 60 mL 0.3% (w/v) PVA solution under continuous stirring at 700 rpm. After 2 hours of continuous stirring, the mixture was transferred to a vacuum chamber and stirred at 300 rpm to further remove DCM. The solidified microparticles were filtered through a 40-μm cell strainer, washed with deionized water, and collected by centrifuging at 500 rpm for 10 minutes. After 3 cycles of washing steps, the microparticles were collected and resuspended in ultrapure water, ready for use.

The HIP method has been applied to further improve the drug loading and drug release for nor-LAAM-MP via a modified process [28,29]. We used three counter-ions for complex formation: oleic acid, sodium docusate, and pamoic sodium monohydrate. In brief, nor-LAAM.HCl and counter-ions were dissolved in deionized water separately, and the counter-ion solution was dropwise added to the nor-LAAM solution under continuous stirring at 300 rpm. For all the counter-ions, a precipitate was observed, confirming a complex formation; **Fig. S1** shows an example of nor-LAAM and pamoic acid complex formation. The resultant complex was centrifuged at 3000 rpm for 15 minutes, and the precipitate was collected and freeze-dried. The polymer and the formed nor-LAAM HIP complex were dissolved in DCM, and then the emulsion solvent evaporation process was followed to prepare the nor-LAAM-MP, as described above.

#### 2.3.1. Characterization of Nor-LAAM-MP

The average particle size of nor-LAAM-MP was analyzed using Malvern Mastersizer 3000. The surface morphology of the lyophilized nor-LAAM-MP was characterized by scanning electron microscope (SEM) using Hitachi SU-70 FE-SEM (Hitachi High-Tech America Inc., USA) after sputter coating. The SEM images were taken under a 15 mm working distance and accelerated voltage of 3 and 5kV.

A certain amount of lyophilized nor-LAAM-MP was weighed and dissolved in 1 mL ACN to measure the drug loading. The drug level was measured by HPLC-UV (Shimadzu Prominence LC system) using a pursuit 5 C18 column, and the mobile phase comprised of acetonitrile and 10 mM KH_2_PO_4_ at pH 4.8 (60:40 *v*/*v*) (flow rate of 1 mL/min). The drug loading (DL) and encapsulation efficiency (EE) were calculated using the following equations:

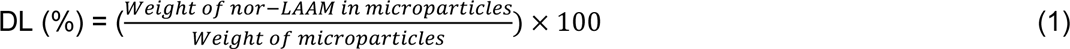

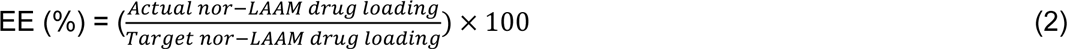

The *in vitro* drug release kinetics of nor-LAAM-MP with and without HIP were evaluated in PBS (pH 7.4) release medium under a sink condition at 37°C on a platform shaker. The nor-LAAM-MP (containing ∼700 μg nor-LAAM) were suspended in 5 mL of PBS in triplicate in scintillation vials. At predetermined time points, 3 mL of the release medium was pipetted from the scintillation vials and replaced with 3 mL of fresh PBS (pH 7.4). As described earlier, the amount of nor-LAAM released from microparticles at each timepoint was then analyzed using HPLC-UV.

### 2.2. Animals

The Virginia Commonwealth University Institutional Animal Care and Use Committee approved the study protocols. For pharmacokinetics studies, New Zealand white rabbits of mixed gender weighed between 2.5 and 2.8 kg were obtained and singly housed with unrestricted access to food and water. For efficacy studies, we obtained adult Sprague Dawley (SD) rats, mixed genders, approximately 11-12 weeks old (250-300 g); each rat was placed in an individual housing with unrestricted access to Teklad Rat Diet food (Envigo) and water. All the animals were obtained from Envigo in Frederick, MD. The vivarium where the animals were housed maintained a controlled environment, including regulated temperature (23 ± 2°C), relative humidity (55 ± 10%), and a 12-hour light/dark cycle.

### 2.4. Pharmacokinetics study in rabbits

Rabbits were randomly divided into three groups, each containing three rabbits: two receiving either nor-LAAM solution intravenously or orally, and the third receiving a single subcutaneous (SQ) injection of nor-LAAM-MP for a month. The nor-LAAM.HCl solution was prepared in sterile water, and microparticles were freshly prepared.

Intravenous (IV) and oral nor-LAAM groups were anesthetized with ketamine (50 mg/kg) and xylazine (5-7 mg/kg) via intramuscular injection before nor-LAAM administration. Five minutes before drug administration, baseline blood plasma was collected from the marginal ear vein. For the oral group, nor-LAAM at 1 mg/kg was administered before the rabbits became unresponsive from anesthesia. Catheters were implanted for both groups, and IV nor-LAAM at 0.1 mg/kg was administered via the catheter. Blood samples were collected at predetermined times: 5, 15, 30 minutes, and 1, 2, 3, 4, 8, 12, 24, 48, 72, 96, and 120 h.

For the group that received nor-LAAM-MP, they were with ketamine (50 mg/kg) and xylazine (5-7 mg/kg) via intramuscular injection, and five minutes before the nor-LAAM-MP administration, baseline blood plasma was collected. After collecting baseline blood plasma, nor-LAAM-MP at a dose of 5 mg/kg was administered, and the blood plasma samples were collected at 2, 4, 8, and 24 h, and at days 4, 8, 15, 22, and 30. The collected blood samples were transferred into EDTA (0.5 M) coated tubes, followed by centrifugation (3000 X g, 3 minutes) to separate plasma. The plasma was stored at −80°C until further analysis (Supplementary materials for detailed bioanalysis). After the study, rabbits were euthanized following intravenous administration of phenobarbital (100 mg/kg).

### 2.5. Efficacy of nor-LAAM on fentanyl-dependent rats

#### 2.5.1. Catheter maintenance and Apparatus

For experimental purposes, the rats underwent surgery to have vascular access ports, and intravenous jugular catheters (Instech, Plymouth Meeting, PA) were implanted, a procedure previously detailed [30]. Subcutaneous administration of ketoprofen (5 mg/kg) was given immediately after surgery, and repeated dosing was done 24 hours later to manage postoperative discomfort. Following surgery, the rats were allowed a five-day recovery period before commencing their fentanyl self-administration training. During fentanyl self-administration training, catheters were flushed with 0.1 mL of gentamicin (4 mg/mL) and heparin saline (30 U/mL). A volume of 0.1 mL methohexital (0.6 mg) was intravenously administered periodically to check catheter patency, a rapid loss of muscle tone as an indicator of catheter patency.

Operant chambers housed within sound-attenuating cubicles for self-administration training had two retractable levers and a 0.1 mL dipper cup for dispensing liquid food. In this study, the liquid food used was a 32% *v*/*v* vanilla-flavored Ensure^TM^ (Abbott Laboratories, Chicago, IL) prepared using tap water obtained from Abbott Laboratories. Each lever in the operant chamber featured three LED lights: yellow (20-s time out), green (Fentanyl presence), and red (food presence).

#### 2.5.4. Drug-vs-Food choice procedure

Once the rats had completed their training in fentanyl and food reinforcement (described in supplementary materials), they were introduced to a drug-vs-food choice protocol following the methodology described in reference [30]. In brief, weekday sessions between 2 to 4 PM were divided into five 20-minute segments (1-5, respectively). Each segment involved different fentanyl doses (0, 0.32, 1, 3.2, and 10 µg/kg/infusion). The liquid food concentration remained consistent in each segment, consisting of 0.1 mL of 32% Ensure. Within each response segment, a non-contingent infusion of the specified fentanyl dose for the upcoming segment was administered, followed by a 2-minute timeout period. Subsequently, a 5-second presentation of liquid food was given, followed by another 2-minute timeout. After the timeout, the choice segment began, with the right and left levers extending. The green light above the right lever indicated the availability of fentanyl, while the red light signaled the availability of liquid food. The session operated on an FR5 schedule, meaning that five consecutive responses on the right lever resulted in IV fentanyl, while five consecutive responses on the left lever led to liquid food presentation. During each segment, rats could fulfill up to ten response requirements between the levers associated with food and fentanyl. The stability of the fentanyl-vs-food choice was considered achieved when the lowest unit dose of fentanyl that maintained over 80% fentanyl choice remained within a 0.5 log unit range over three consecutive days. Baseline values for analysis were determined by averaging the data collected from Monday to Friday. After the drug-vs-food training reached stability, an extended fentanyl access period was initiated, lasting from 6 PM to 6 AM for two weeks (Sunday to Friday). During this period, rats could self-administer 3.2 µg/kg/infusion of fentanyl under an FR5 schedule with a 10-second timeout period. The day following the first extended fentanyl access session, two groups were treated with either high-dose (Nor-LAAM-MP-70 mg/kg) or low-dose (Nor-LAAM-MP-20 mg/kg) doses of nor-LAAM-MP and one group received vehicle saline. Before the start of each daily session, the rats’ body weights were recorded, and a 30-second observation period was conducted to assess the presence of nine somatic withdrawal signs. These signs included paw tremors, ptosis, diarrhea, wet dog shake, yawn, mastication, piloerection, teeth chatter, and eye twitch.

### 2.6. Histopathology evaluation

At the endpoint of the efficacy study, rats were euthanized using CO2 euthanasia followed by cervical dislocation. We collected rat tissues, including the skin tissue at the injection site, liver, kidney, spleen, and heart. All mentioned tissues were fixed in a 10% formalin solution for a day and then transferred to 70% ethanol. The tissues were embedded in paraffin, then sliced into 5 μm sections and stained with hematoxylin and eosin (H&E) for histology evaluation. A blinded pathologist evaluated the H&E-stained slides for any signs of tissue damage, inflammation, or other histological changes. Images were obtained using phenochart, a tissue imaging software penoimager (AKOYA Biosciences).

### 2.7. Data analysis

For analyzing pharmacokinetics studies, we used one-way ANOVA followed by the Dunnett test to compare the bioavailability data between oral and SQ administration of nor-LAAM. For analysis of the efficacy study, the dependent measures included the percentage of fentanyl choices made, the number of choices completed per segment, the presence of withdrawal signs, changes in body weight as a percentage, and the number of fentanyl injections earned during the extended access sessions. We used both one-way and two-way repeated measures ANOVAs for analysis. We employed the non-parametric Friedman test, followed by Dunn’s multiple comparisons for post-hoc analysis. Data were considered significant if the p-value was < 0.05). All data were analyzed using GraphPad Prism.

## 3. Results and Discussion

### 3.1. Formulation and characterization of nor-LAAM Microparticles (nor-LAAM-MP)

Our initial experiments with a conventional emulsification method using PLGA_18kDa_ only resulted in 3 wt.% DL when the target DL was 20 wt%. This limited DL may be attributed to the water-soluble nature of nor-LAAM.HCl, which makes encapsulation challenging. We proceeded with a new strategy, converting the water-soluble nor-LAAM.HCl salt into the less water-soluble nor-LAAM free base using triethylamine (TEA) during the emulsification process. Although the DL increased significantly to 13 wt.%, an undesired burst release (nearly 40% within 24 hours) was observed, which might be due to a large amount of drug that is located on the microparticle surface [31]. Apart from the significant initial burst release, the nor-LAAM free base formulation had a short *in vitro* release profile (∼70% in 7 days), which rendered this formulation unsuitable for *in vivo* applications (**Fig. S2**). It was shown that tertiary amine drugs, such as risperidone and olanzapine, can cause basic catalysis that can lead to polymer degradation due to polymer hydrolysis because of the basic environment [32,33]; it is possible that nor-LAAM, containing a tertiary amine with a pKa of 10.34, could drastically hydrolyze PLGA, leading to accelerated release of the drug from microparticles.

We adopted a HIP[34–36] by forming a lipophilic HIP complex between nor-LAAM.HCL and counterions that could better partition into the hydrophobic core of the polymer matrix during the encapsulation process. In this study, three different formulations were developed: nor-LAAM/pamoic acid HIP (F1), nor-LAAM/oleic acid HIP (F2), and nor-LAAM/docusate HIP (F3) (**Fig. 1**). The formed nor-LAAM-loaded PLGA_18kDa_ microparticles had sizes of 25 µm, 18 µm, and 20 µm, respectively, but with notable surface micropores on F1 formulation compared to F2 and F3 formulations (**Fig. 2A-C**). The F2 formulation with oleic acid HIP exhibited electric charge buildup during SEM imaging, which could have been from the poor platinum sputter coating because of the oily oleic acid located on the particle surface [37,38]. For all three formulations (nor-LAAM-MP-F1, F2, F3), we achieved high DL (∼13 wt.%) when the target DL was set at 20% and provided sustained release of nor-LAAM *in vitro* conditions (**Fig. 2D**). Nor-LAAM-MP-F2 and F3 formulations showed a biphasic *in vitro* drug release profile with an initial rapid release, followed by a slow release. The nor-LAAM-MP-F1 formulation exhibited a monophasic drug release profile with a steady in vitro drug release. The release kinetics may be attributed to the relatively larger molecular size complex with a smaller diffusion coefficient of the pamoate HIP (total molecular weight 1184.18 g/mol) compared to oleic acid HIP (total molecular weight 659.39 g/mol) and docusate HIP (total molecular weight 820.49 g/mol). The molecular weight of the formed complex was calculated after revealing that the mole ratio between nor-LAAM and pamoic acid was 2:1 and 1:1 for the oleic acid and docusate. The larger HIP drug complex potentially led to slower drug diffusion and release through the PLGA polymer matrix [35].

**Fig. 1.**
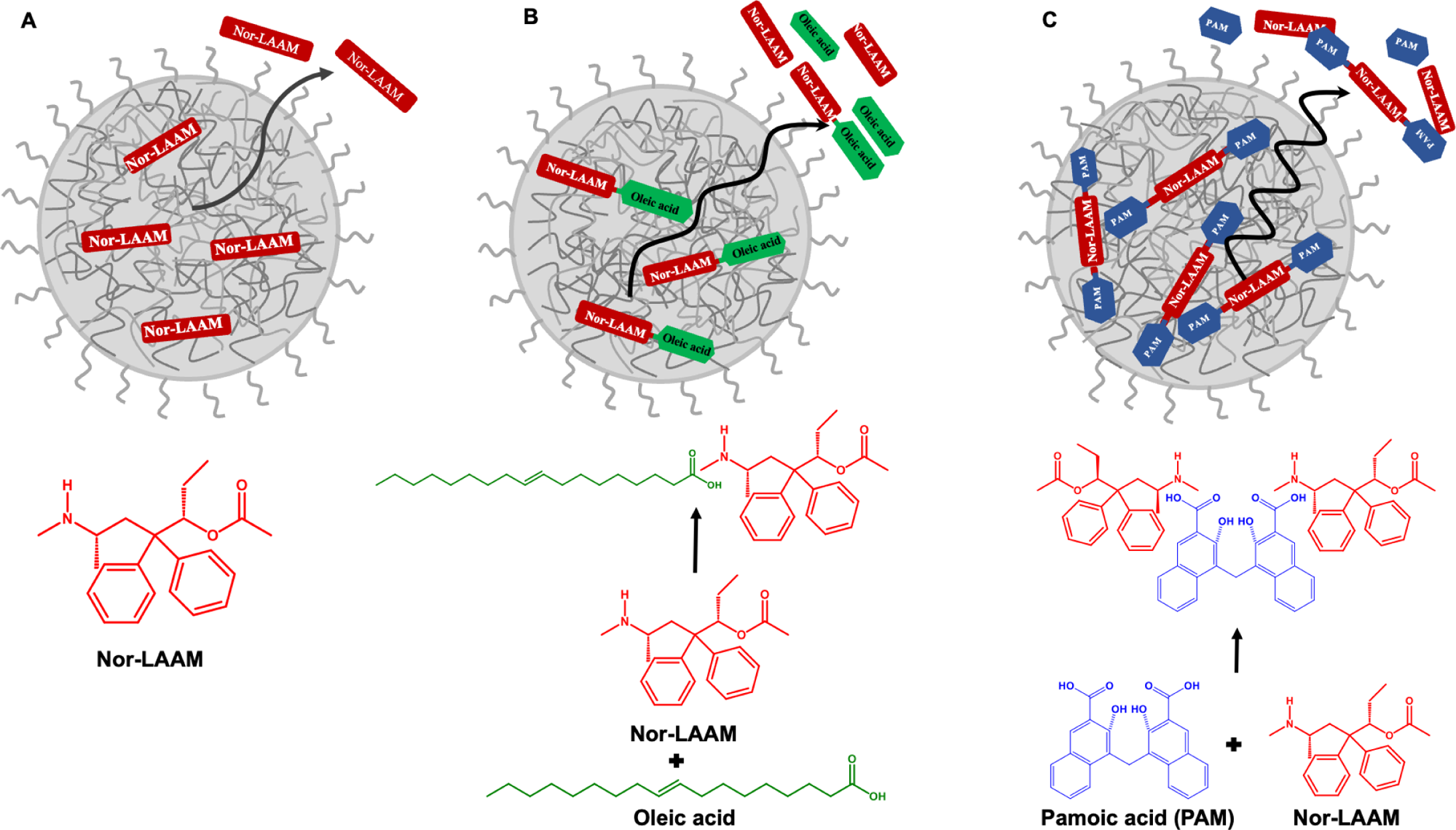
A schematic showing the HIP formation. **(A)** nor-LAAM only, **(B)** nor-LAAM with oleic acid, **(C)** nor-LAAM with pamoic acid.

**Fig. 2.**
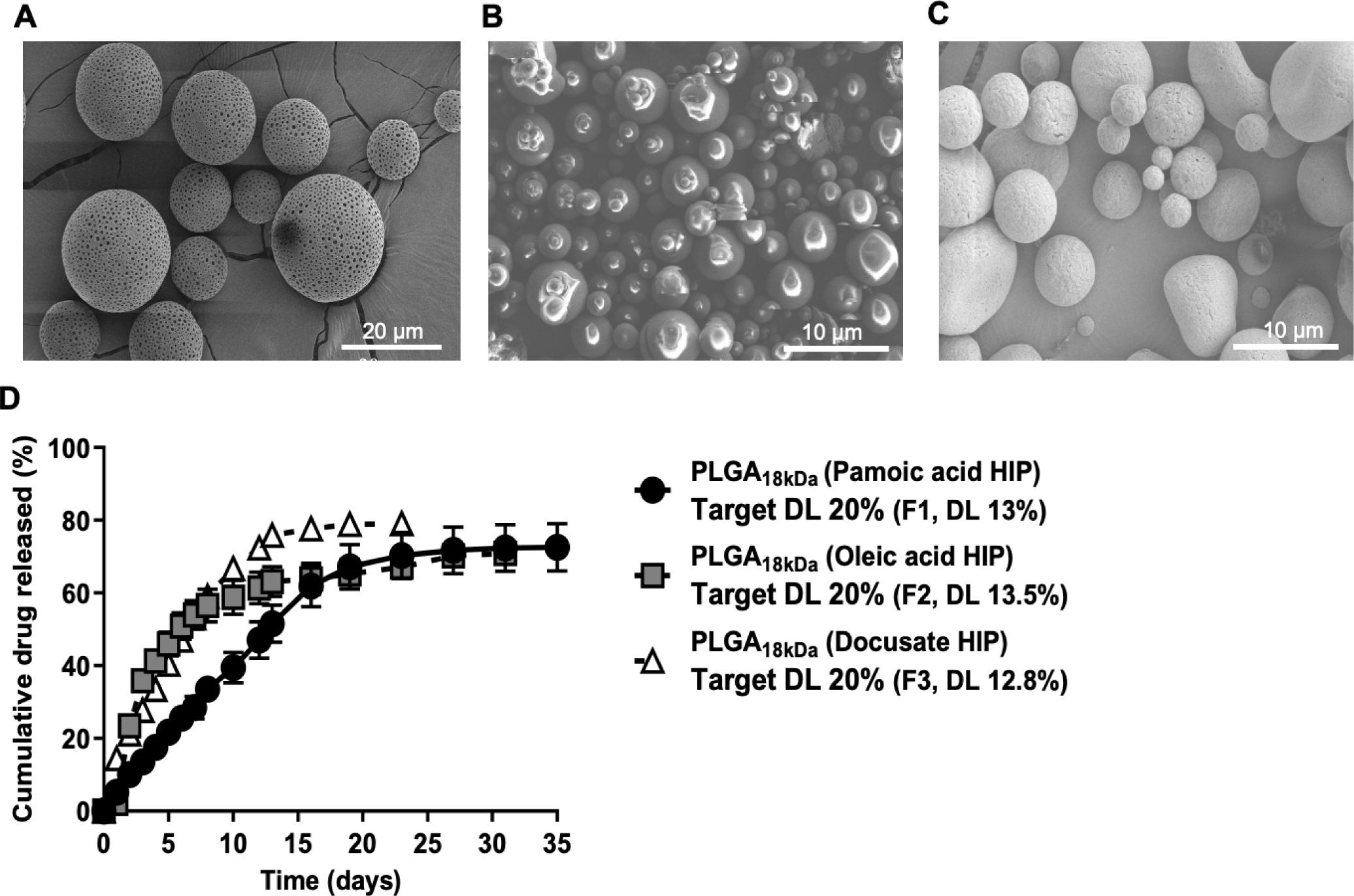
Representative SEM image of PLGA_18kDa_ nor-LAAM-MP **(A)** F1, **(B)** F2, **(C)** F3. **(D)** Nor-LAAM-MP prepared by HIP with different counterions pamoate (F1), oleic acid (F2), and docusate (F3) exhibited high drug loading ∼13 wt.% and provided sustained release without obvious burst *in vitro*. (n=3, mean ± SEM).

To further optimize nor-LAAM-MP suitable for *in vivo*, we explored the use of poly (ethylene glycol) (PEG)-coated particles since PEG polymer has been shown to mitigate inflammation risk associated with locally injected polymeric nanoparticles and microparticles [39,40]. The nor-LAAM-MP with pamoate HIP was selected because of its high drug loading and more steady drug release. Pamoic acid has proven to be safe, with no systemic toxicity, and has been extensively used in FDA-approved pharmaceutical products [41]. For sustained-release therapeutic effects of OUD, we developed PEGylated PLGA_18kDa_/PLGA_45kDa_-PEG_5kDa_ (1:4) pamoate HIP MP. The target DL was F4: 20%, F5: 30%, and F6: 40%, respectively. For all the PEGylated nor-LAAM-MP (F4, F5, and F6), we achieved a high DL of 11 wt.% (F4), 14 wt.% (F5) and 18 wt.% (F6), and the mole ratio between nor-LAAM and pamoic acid remained approximately 2:1. The resulting PEGylated nor-LAAM-MP-(F4, F5, and F6) had an average particle size of 20 µm, 24 µm, and 26 µm, respectively, and smooth surfaces without obvious pores (**Fig. 3A, B, C**). For sustained drug release, we tested the release mechanism of nor-LAAM and pamoic acid, and all the PEGylated nor-LAAM-MP formulations exhibited a sustained *in vitro* nor-LAAM release for over a month without obvious initial burst release. Pamoic acid exhibited similar release trends (**Fig. 3D and Fig. S3**). These results showed that PEGylation did not compromise the nor-LAAM-MP drug loading and sustained drug release profiles, as we observed in the previous F1 formulation.

**Fig. 3.**
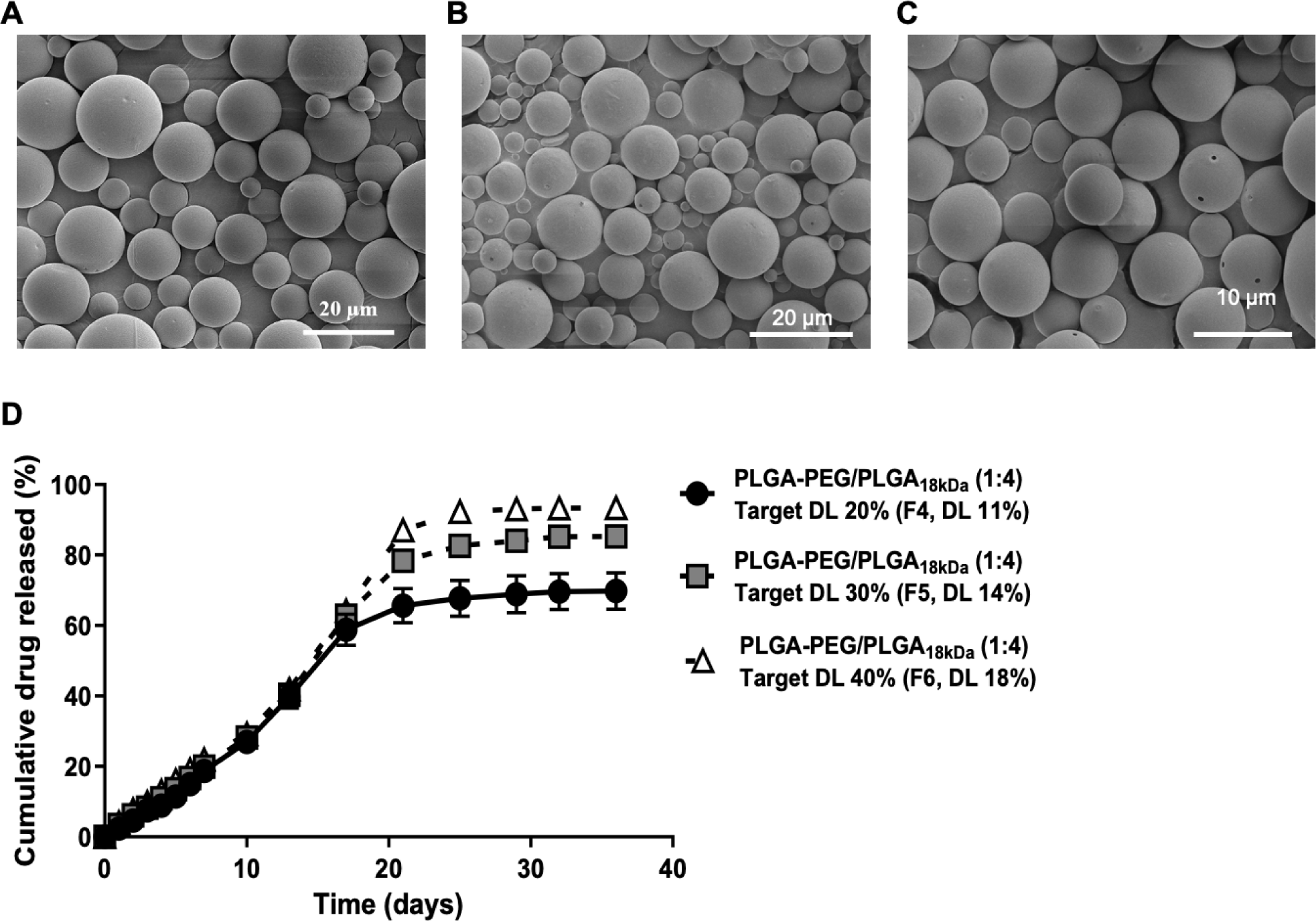
PEGylated nor-LAAM-MPs prepared by pamoate HIP strategy with high target drug loading. Representative SEM image of PLGA_45kDa_-PEG_5kDa_/PLGA_18kDa_ (1:4) **(A)** F4, **(B)** F5**, (C)** F6. **(D)** In vitro drug release profile for PEGylated nor-LAAM-MPs (F4, F5, and F6), all formulations exhibited high drug loading and provided sustained release without obvious bursts in vitro. Mean ± SEM (n=3).

Furthermore, increasing the target DL of no-LAAM did not improve the encapsulation efficiency (**Fig. 3D**). The outcome may be attributed to the saturation of nor-LAAM-HIP within MP, where further drug encapsulation became unattainable. This suggested that the drug may have reached its solubility limit within the MP, resulting in no additional drug being encapsulated [42]. Overall, we achieved high DL and sustained drug release without apparent initial burst release for the nor-LAAM-MP developed using the HIP strategy.

### 3.2. In vitro drug release kinetics toward achieving sustained drug release

The physicochemical properties of polymers and the interaction between nor-LAAM-HIP and polymers could be the key factors for controlling drug release kinetics. It is currently known that an extended *in vitro* drug release at a constant rate (zero-order) without any initial burst or delay (monophasic) can achieve an ideal *in vivo* pharmacokinetic profile [43]. We developed several microparticles to optimize drug release kinetics using different Mw of PLGA (either carboxyl or ester terminal group) and co-polymers with PLGA-PEG. As shown in **Fig. 4A**, among all the formulations, MP with the lowest Mw polymer (PLGA_5kDa_) exhibited an intense initial burst release of nor-LAAM (more than 10% within 24 hours). This initial burst release could be related to the drug trapped near the microparticles’ surface, leading to a large initial drug concentration gradient between the system and the medium. Increasing the Mw of polymers reduced the initial burst release of nor-LAAM, which concluded a negative correlation between initial polymeric Mw and initial burst release. To note, F1 (PLGA_18kDa_) showed over 2-fold higher drug release within 24 hours than F4 (PLGA_45kDa_-PEG_5kDa_/PLGA_18kDa_ 1:4), indicating the porous surface of F1 (**Fig. 2A**) exhibited faster water accessibility to remove drug molecules near the particle surface compared to nonporous F4 (**Fig. 3A**). However, the porous vs. nonporous systems did not affect the drug release kinetics at the second phase between F1 and F4, which contained similar polymeric Mw (_18kDa_) (**Fig. 2D, 4B**). It indicates that the PLGA can effectively trap lipophilic pamoate nor-LAAM and is likely liberated by consistent diffusion through the polymeric barriers during the second release phase (2-17 days) (**Fig. 4B**). Surprisingly, an initial ‘lag phase’ followed by a rapid release of the drug was observed when microparticles F7 (carboxyl terminated PLGA) and F8 (ester terminated PLGA) contained large polymeric Mw (PLGA_45kDa_-PEG_5kDa_/PLGA_54kDa_ 1:4) (**Fig. S4**).

**Fig. 4.**
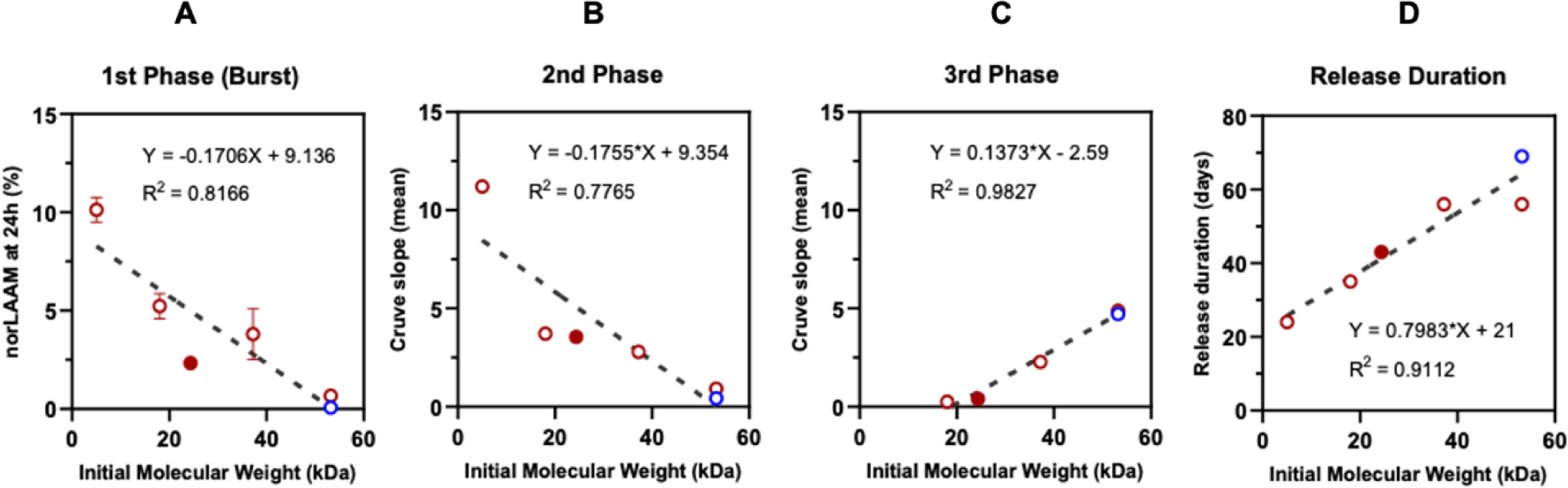
Cumulative nor-LAAM release from microparticles *in vitro* (PBS 0.1 M, pH 7.4). Correlations between polymers’ initial Mw and the different phases of release. (**A**) Percentage of total nor-LAAM released during the burst phase (1^st^ phase) after 24h of incubation. (**B**) The slope of the linear fit during the 2^nd^ phase of nor-LAAM release (day 2 to day 17). (**C**) The slope of the linear fit during the 3^rd^ phase of nor-LAAM release (day 20 to day 30). (**D**) Release duration in days for the microparticles (F1, F4, F7, and F8). The blue circle is the ester terminal group on PLGA. The solid spot is the formulation for *in vivo* studies. Each point is a mean ± SEM (n=3).

Given that, the large amount of pamoate nor-LAAM-HIP started to diffuse and liberate from the core of microparticles as soon as the polymer erosion occurred rapidly after 2 weeks. A strong positive correlation (0.798) and good linearity with correlation coefficients 0.91 were then found between initial polymeric Mw and release duration (**Fig. 4D**). Remarkably, the release duration of F8 was around 2 weeks longer than F7. The results align with the previous finding, showing carboxyl terminal groups could increase acidic conditions inside the microparticles, leading to autocatalytic acceleration of central polymer degradation [44]. Since nor-LAAM (an active metabolite of LAAM) shows much more potency than the parent [45], the initial burst release could cause a safety concern if the systemic drug levels reach above the toxicity threshold. Furthermore, an initial ‘lag phase’ followed by a rapid drug release may require additional prescription (e.g., oral dosage form) to compensate for this lag period, causing inconvenience. After thoroughly analyzing the release profiles exhibited by these formulations, the data presented supported the selection of nor-LAAM-MP-F4 to be tested in the following *in vivo* studies. Notably, nor-LAAM-MP-F4 demonstrated a sustained release profile of nor-LAAM, characterized by a consistent rate of delivery without any initial burst or delays.

### 3.3. In vivo pharmacokinetics of nor-LAAM loaded PLGA microparticles

In the rabbit pharmacokinetic (PK) studies, blood samples were collected at predetermined time intervals to evaluate the systemic exposure of our optimized nor-LAAM microparticles, as illustrated in **Fig. 5A**. Plasma PK concentration-time profiles were illustrated in **Fig. 5B-D**, and PK parameters were summarized in **Table 2**. The average area under the plasma concentration-time curve from 0 to infinity (AUC_inf_) values calculated following oral solution, IV solution, and SQ microparticle administration of 0.1, 1, and 5 mg/kg of nor-LAAM were 124.5, 343.8, and 5110.7 h×ng/mL, respectively. The mean maximum plasma concentration (C_max_) values obtained following oral solution and SQ microparticle administration were 37.0 and 64.5 ng/mL, respectively. The mean time to reach C_max_ (t_max_) obtained after oral solution and SQ microparticle administration were 0.25 and 8 h, respectively. The absolute oral bioavailability of nor-LAAM was 27.6 ± 22.9 %; more importantly, F4 formulation following SQ administration exhibited a nearly 3-fold higher absolute bioavailability (**Table 2**), based on the dose-normalized AUC_inf_, compared to oral administration of nor-LAAM solution in rabbits. The short elimination half-life of 3.3 hours following IV injection of nor-LAAM solution may be attributed to its relatively high clearance of nor-LAAM.

**Fig. 5.**
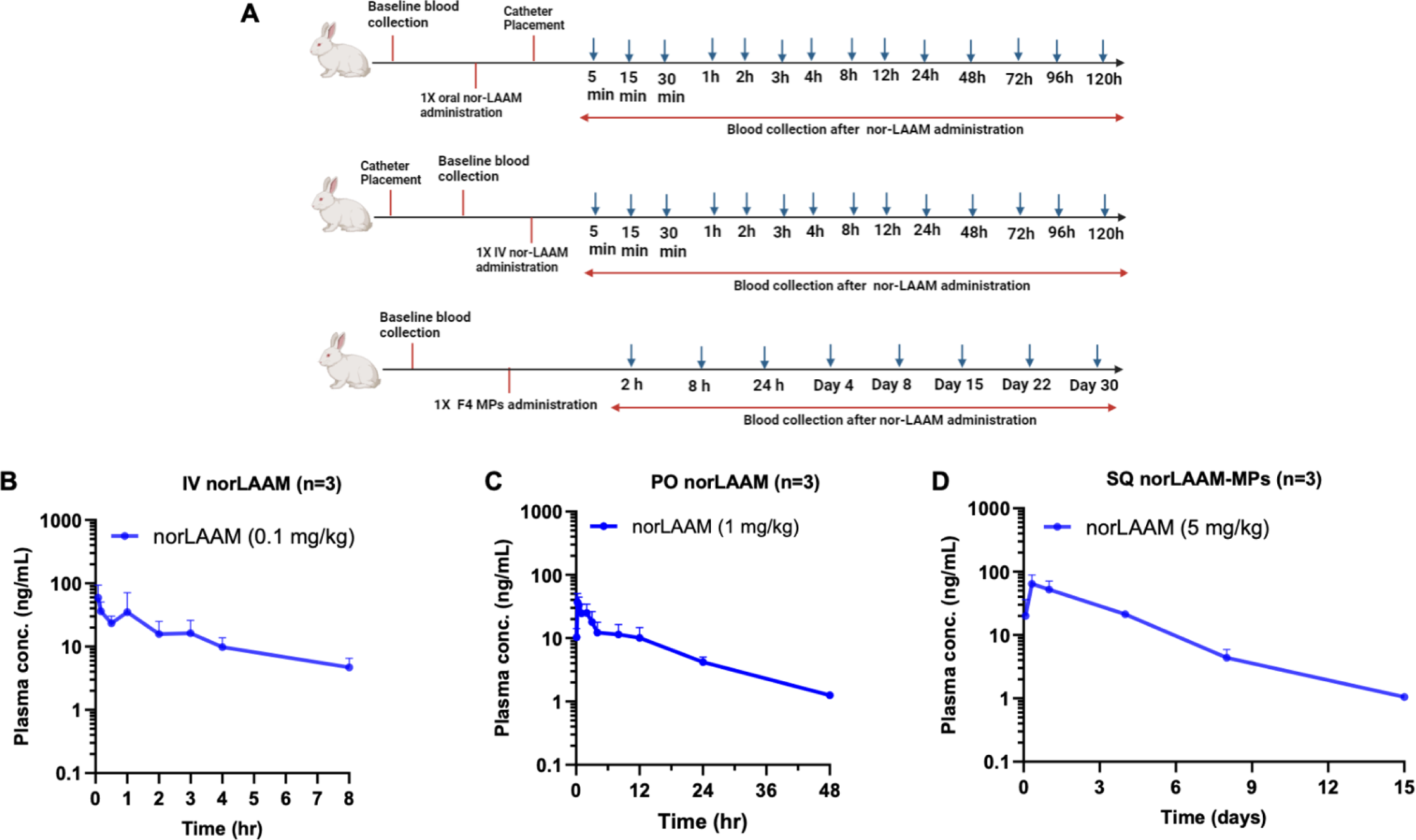
Comparative plasma pharmacokinetic profiles of nor-LAAM in rabbits following different routes of administration. **(A)** The timeline for the PK study with nor-LAAM oral and IV and subcutaneous administration. **(B)** Single intravenous bolus administration (IV) of nor-LAAM (0.1 mg/kg). **(C)** Single oral administration (PO) of nor-LAAM (1 mg/kg). **(D)** Single subcutaneous injection (SQ) of nor-LAAM-MP-F4 (5 mg/kg). Data are presented as mean ± SD.

**Table 2.** Plasma pharmacokinetic parameters of norLAAM following administration of nor-LAAM to rabbits (mean ± SD, n = 3).

| Formulation | Nor-LAAM solution | No-LAAM solution | Nor-LAAM-MPs |
| --- | --- | --- | --- |
| Route of administration | IV (0.1 mg/kg) | PO (1mg/kg) | SQ (5mg/kg) |
| $AUC_{inf}$ (h×ng/mL) | 124.5 ± 39.6 | 343.8 ± 90.7 | 5110.7 ± 1082.2 |
| $AUC_{0\rightarrow t}$ (h×ng/mL) | 96.5 ± 17.0 | 299.7 ± 128.8 | 4932.5 ± 932.6 |
| $C_{max}$ or $C_0$ (ng/mL) | 54.0 ± 26.2 | 37.0 ± 13.7 | 64.5 ± 24.3 |
| **t<sub>max</sub> (h)** | — | 0.25 | 8 |
| **t<sub>1/2</sub> (h)** | 3.9 ± 1.7 | 13.7 ± 3.6 | 55.3 ± 9.3 |
| **CL (mL/h/kg)** | 866.5 ± 330.1 | — | — |
| **V<sub>ss</sub> (mL/kg)** | 4726.5 ± 18996.3 | — | — |
| **F (%)** | — | 27.6 ± 22.9 | 82.1 ± 54.7 * |

AUC<sub>inf</sub>, area under the curve from time zero to infinity; AUC<sub>0→t</sub>, area under the curve from time zero to the last sampling time point; C<sub>max</sub>, maximum observed concentration; t<sub>1/2</sub>, half-life. Significant differences between PO and SQ: \*p ≤ 0.05.

Nor-LAAM concentration was quantified in the rabbit plasma for up to 2 weeks following SQ microparticle administration; after two weeks, nor-LAAM was below the limit of quantification (LOQ: 0.2 ng/mL) (**Fig. 6D**). The *in vitro* release results showed that over 70% of nor-LAAM was released consistently within the first 2 weeks, compared to a further ∼20% over the following 2 weeks. This significant reduction in the *in vitro* nor-LAAM release after the second week can explain the undetectable systemic drug levels after day 15 post-administration in rabbits. Comparable PK data were also evident in rat pilot efficacy studies (**Fig. S5**), where after oral nor-LAAM administration (5.6 mg/kg, n=3) remained detected for up to 72 hours, and after no-LAAM-MP (70 mg/kg, n=2) nor-LAAM was detected on day 7 and day 18. The *in vivo* degradation of PLGA did not correlate with the *in vitro*, and that could have been due to the cellular infiltration in the SQ injection site (**Fig. 8**), where macrophages can create an acidic microenvironment to accelerate polymeric digestion [46]. The one-month PK study in rabbits involving nor-LAAM-MP-F4 (5 mg/kg) showed no lethality or apparent changes in physical behaviors, including body weight change (**Fig. S6),** food and water intake, and diarrhea, indicating the great safety and tolerability of our novel microparticle formulation. These results collectively highlighted the superior bioavailability achieved following a single SQ injection of nor-LAAM-MP, compared to oral nor-LAAM administration.

**Fig. 6.**
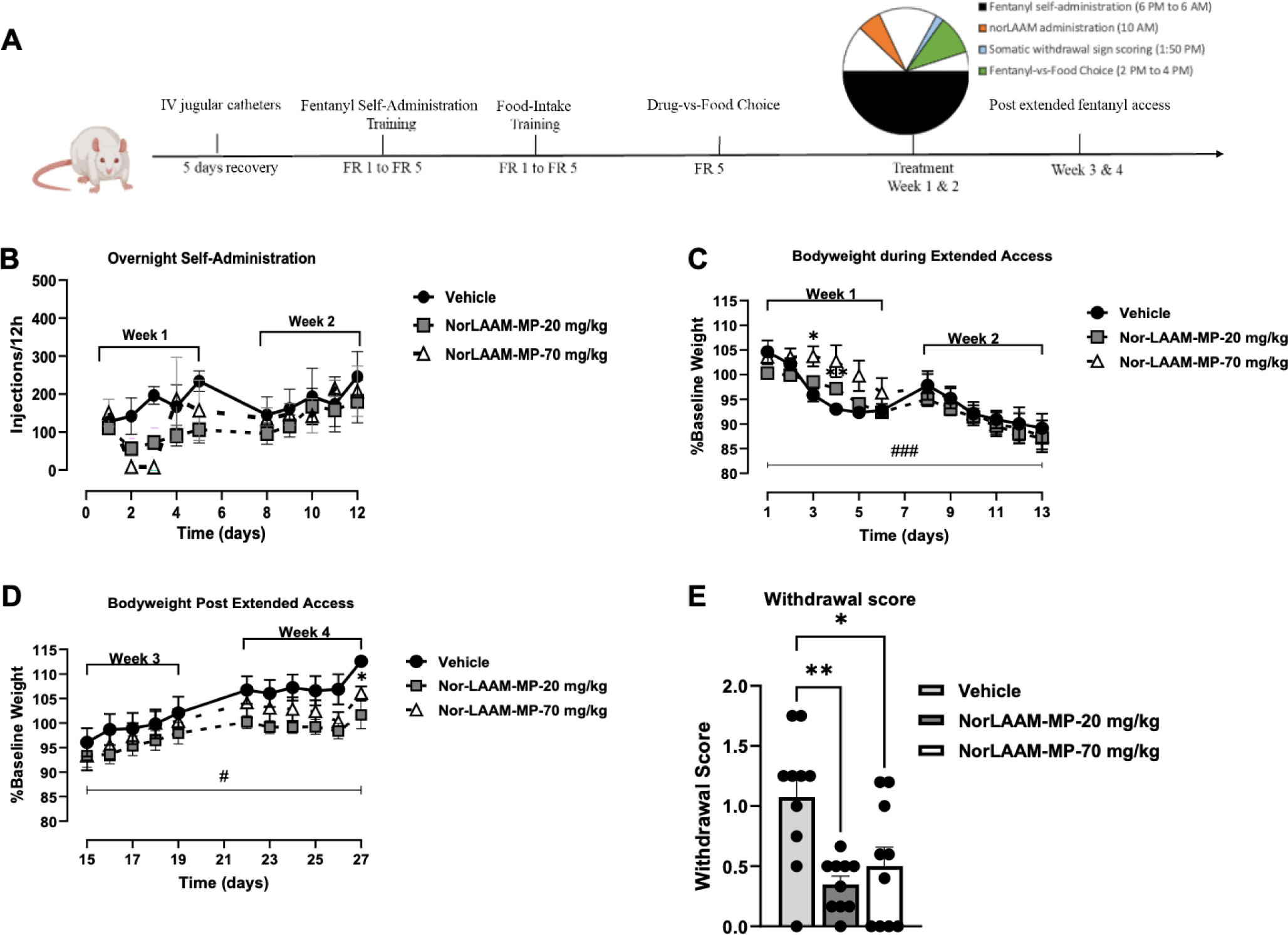
Extended fentanyl self-administration (6 PM - 6 AM) effects on somatic withdrawal sign expression and body weight in the vehicle-treated group (n = 4), nor-LAAM-MP-20 mg/kg treated group (n = 6), and nor-LAAM-MP-70 mg/kg treated group (n = 5). (A) Timeline of the study. (B) Number of fentanyl injections (3.2 mg/kg/inj) self-administered during a 12-hour session. (C) Changes in body weight were expressed as a percentage of baseline during the extended access (weeks 1 and 2) and (D) Post-extended access (weeks 3 and 4). (E) The withdrawal score was observed after each extended access self-administration session. Data are represented as mean ± SEM. Significant differences between treatment group and vehicle: *p ≤ 0.05, ** p ≤ 0.01; # symbolizes time and bodyweight interaction (Two-Way ANOVA).

### 3.4. Nor-LAAM Efficacy Study on fentanyl-dependent rats

To assess the therapeutic potential of nor-LAAM as a novel OUD pharmacotherapy, we employed a preclinical approach involving fentanyl self-administration to induce opioid dependence and somatic withdrawal in SD rats. Drug self-administration is a well-established methodology that has previously demonstrated sensitivity to clinically used OUD medications such as methadone and buprenorphine on opioid choice in both opioid-dependent rats and rhesus monkeys [23,24]. Given the absence of prior data regarding the effectiveness of nor-LAAM, our initial investigation sought to examine the influence of orally administered nor-LAAM solution in attenuating fentanyl choice among fentanyl-dependent rats. Additionally, we conducted an extended assessment of the effects of sustained-release nor-LAAM-MP-F4. **Figure 6A** provides an overview of the study timeline. Before nor-LAAM treatment, all rats chose food over fentanyl when no or small units of fentanyl doses were available (0, 0.32, 1 µg/kg/injection). As the unit fentanyl dose increased, rats reallocated their behavior away from food and towards fentanyl such that at the largest fentanyl dose (10 mg/kg/infusion), rats were exclusively choosing fentanyl. Once the rats began extended fentanyl sessions in addition to the 2-h fentanyl-food choice sessions, an oral 5.6 mg/kg nor-LAAM solution was administered *via* oral gavage three times weekly over two weeks. Oral administration of nor-LAAM solution significantly reduced fentanyl choice and shifted preference to food in rats at fentanyl doses of 3.2 and 10 µg/kg/infusion compared to the vehicle-treated group.

Additionally, oral nor-LAAM significantly decreased somatic withdrawal signs (\**p* < 0.05) (**Fig. S7**); we demonstrated that the nor-LAAM oral solution (5.6 mg/kg) administered 3 times per week (M-W-F) effectively reduced fentanyl choice like 5.6 mg/kg methadone solution administered 3 times per day (**Fig. S8**). After demonstrating the effectiveness of oral nor-LAAM solution (5.6 mg/kg) in decreasing fentanyl choice in opioid-dependent rats, the efficacy of a single subcutaneous dose of nor-LAAM-MP-F4 was investigated for four weeks. Two doses were examined: a low dose (nor-LAAM-MP-20 mg/kg) and a high dose (Nor-LAAM-MP-70 mg/kg) with a control group receiving saline vehicle treatment. The initial first week of extended fentanyl access showed a trend towards a decrease in fentanyl choice in both the low-dose and high-dose nor-LAAM-MP treatment groups. However, this reduction did not reach statistical significance. By the second week, fentanyl choice levels had equalized among all treated groups, as depicted in **Fig. 6B**. Body weight decreased over time across the two-week extended fentanyl self-administration period with an F-statistic calculated to be F_22, 143_ = 3.011, *p* < 0.001, as indicated in **Fig. 6C** with the vehicle-treated group showing rapid weight decrease compared to the other treatment groups, confirming the development of opioid dependence. In the post-extended fentanyl access period (weeks 3 and 4), a recovery in body weight was observed over time (F_20, 114_ = 1.785, p = 0.030), as shown in **Fig. 6D**. Furthermore, the nor-LAAM-MP-20 mg/kg and 70 mg/kg doses significantly decreased somatic withdrawal signs compared to the vehicle (Friedman test, P < 0.007, n = 6 and Friedman test P < 0.028, n = 5), as illustrated in **Fig. 6E**.

There were no variations in fentanyl response observed in baseline fentanyl choice across all treatment groups, as depicted (**Fig. S9**). The vehicle-treated group exhibited an increase in fentanyl choice during both weeks 1 and 2 and weeks 3 and 4. There was an interaction between fentanyl dose and time-extended access (F_1.815, 5.446_ = 9.424, p = 0.0180); however, the post-hoc analysis failed to detect any significant differences (**Fig. 7A-B, S10 A-B**). The nor-LAAM-MP-20 mg/kg treated group exhibited a similar trend to the vehicle-treated group, with no observable reduction in fentanyl choice during weeks 1, 2, 3, and 4 (**Fig 7C-D, S10 C-D)**. Conversely, there was a significant decrease in fentanyl choice during weeks 1 and 2 after high nor-LAAM dose treatment (F_2.141, 8.565_ = 10.77, p = 0.004) (**Fig. 7E-F**). This trend correlates with the findings from our PK studies, wherein a high concentration of nor-LAAM was detected within 15 days. While a trend towards decreased fentanyl choice was also observed during weeks 3 and 4, no statistical significance was detected (**Fig. S10 E-F**). Notably, the study demonstrated that a single subcutaneous injection of nor-LAAM-MP-70 mg/kg can achieve a significant reduction in fentanyl choice and somatic withdrawal symptoms consistent with the efficacy of oral nor-LAAM solution (three times/week).

**Fig. 7.**
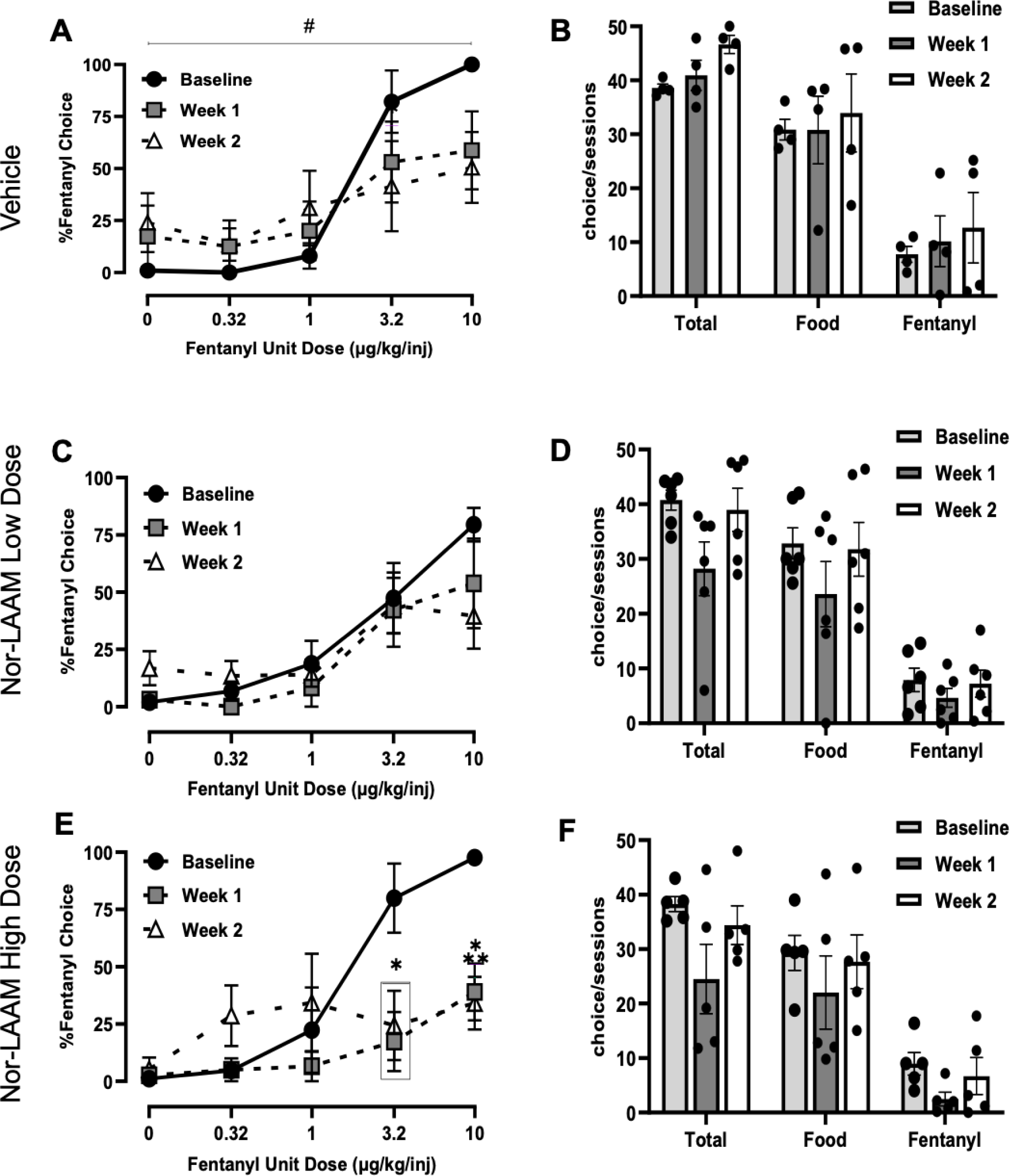
Effects of a single subcutaneous injection of vehicle (n = 4), nor-LAAM-MP-20 mg/kg (n = 6), and nor-LAAM-MP-70 mg/kg (n = 5) on fentanyl-vs.-food choice in opioid-dependent rats. (**A, C, E**) treatment effect on percent fentanyl choice in weeks 1 and 2. (**B, D, F**) Number of fentanyl-vs-food choices completed per session in weeks 1 and 2. Data are represented as mean ± SEM. Significant differences between treatment group and vehicle: *p ≤ 0.05, ** p ≤ 0.01; # symbolizes a significant fentanyl unit dose and time interaction (Two-Way ANOVA).

Upon reaching 30 days of the study, various tissues, including those of the spleen, heart, kidney, and liver, were collected. Histopathologic evaluations showed no evidence of toxic effects in these tissues 30 days post-administration of nor-LAAM-MP. In addition, microparticle depots remained at the injection site and were evident in both the nor-LAAM-MP-20 mg/kg and nor-LAAM-MP-70 mg/kg groups. The microparticle depots (yellow arrows) elicited a robust tissue response, typically seen in the presence of a foreign body and consisting of an almost encapsulating dense lymphohistiocytic infiltrate (green arrows). The spleen tissues showed the usual red and white pulp. The myocardium had no fibrosis, and the cardiac myocytes displayed the usual cytoarchitecture without hypertrophy. Renal tissues revealed pristine glomerular and tubular structures without sclerosis or acute necrosis. The hepatic lobules did not show signs of injury or increased inflammation (**Fig. 8**).

**Fig. 8.**
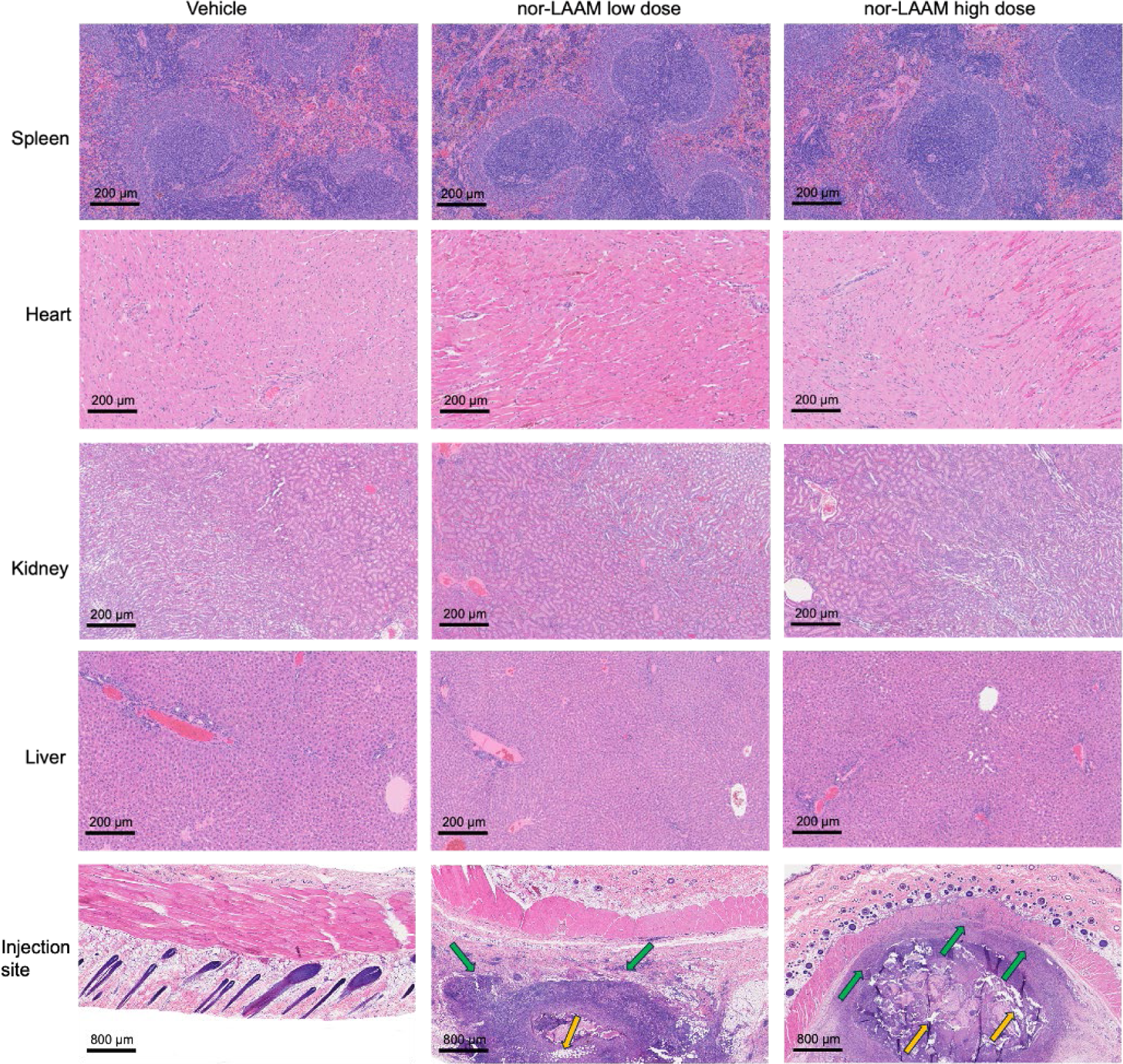
Representative hematoxylin and eosin-stained sections of collected tissues (spleen, heart, kidney, liver, skin injection site) from animals 30 days post-subcutaneous injection of the vehicle and nor-LAAM-MP.

## 4. Conclusion

The current research aimed to address the challenges in treating OUD by studying the nor-LAAM effect in reducing fentanyl choice and alleviating withdrawal symptoms in fentanyl-dependent rats. The current OUD treatment has been shown to be effective; However, some limitations associated with the current treatment are frequent dosing, poor patient retention, poor efficacy, and safety concerns. To address some of these concerns, we successfully developed nor-LAAM-loaded PLGA microparticles that could potentially address some of the limitations mentioned earlier. The development of a lead nor-LAAM-MP-F4 formulation using the HIP technique, when tested *in vivo*, showed that a single SQ injection of microparticles had better bioavailability than an oral nor-LAAM solution in rabbits. Moreover, both nor-LAAM-MP-F4 and nor-LAAM oral solution showed a significant reduction in fentanyl choice and withdrawal symptoms in fentanyl-dependent rats. It suggests the nor-LAAM-MP formulation could be a more effective and long-acting systemic delivery strategy for treating OUD.

Although the findings in the research paper showed some promise for nor-LAAM-MP to be a potential therapy for OUD, it’s crucial to acknowledge that this study had its constraints that could have influenced the data presented. Some key limitations of these studies include (1) failure to achieve 100% drug release for the formulation, which could be from different factors such as losing the drug during the drug release sampling or drug binding to the column during analytical methods. (2) different release kinetics *in vitro* and *in vivo*, and (3) underpowered efficacy studies. We recognize these limitations, which will be addressed in future formulation development and efficacy studies.

## ACKNOWLEDGEMENT

This work was partially funded by grants from the National Institutes of Health (NIH) (UG3DA048768, P30DA033934) and VCU’s Quest Project Award. We acknowledge the Drug Supply Program of the National Institute of Drug Abuse (NIDA) for providing the experimental drug nor-LAAM.HCl. We acknowledge the VCU Tissue and Data Acquisition and Analysis Core Laboratory for the histological analysis. We also acknowledge the NIDA’s T32 graduate training grant (T32DA007027) to D.I. for her PhD study.

## AUTHOR CONTRIBUTION

**Diane Ingabire/Chaolong Qin**: Conceptualization, Data curation, Formal analysis, Investigation, Methodology, Software, Writing – original draft. **Tuo Meng/Aji Alex Moothendathu Raynold**: Conceptualization, Data curation, Formal analysis, Investigation, Methodology, Writing – review & editing. **Hadi Sudarjat**: Investigation, Writing – review & editing. **Andrew Townsend**: Data curation, investigation, Methodology, Software. **Rudra Pangeni/Sagun Poudel**: Writing – review & editing. **Michelle Arriaga:** Methodology**. Long Zhao**: Methodology; Writing – review & editing. **Woon Chow**: Visualization, Writing – review & editing. **Matthew Banks**: Conceptualization, Funding acquisition, Project administration, Supervision, Writing – review & editing. **Qingguo Xu**: Conceptualization, Funding acquisition, Project administration, Supervision, Writing – review & editing.

## DECLARATION OF INTEREST

The nor-LAAM-MP technology described in this publication has been filed as a patent application through Virginia Commonwealth University (VCU). Qingguo Xu, Matthew Banks, Diane Ingabire, Chaolong Qin, Tuo Meng, and Aji Alex Moothedathu Raynold are the co-inventors who are subject to certain rules and restrictions under VCU policy. The terms of this arrangement are being managed by VCU in accordance with its conflict-of-interest policies.

## DECLARATION OF USE GENERATIVE AI AND AI-ASSISTED TECHNOLOGIES

During the preparation of this work, the author(s) used ChatGPT to improve language since English is not the author(s) first language. After using this tool/service, the author(s) reviewed and edited the content as needed and take(s) full responsibility for the content of the publication.

