## Supplementary materials for "Nor-LAAM loaded PLGA Microparticles for Treating Opioid Use Disorder"

### 1. Methods

#### *1.1* ***Bioanalysis of nor-LAAM in plasma***

The concentrations of nor-LAAM in plasma were determined using a Liquid Chromatography Mass Spectrometer (LC-MS-2020) (Shimadzu Corporation). Briefly, a calibration (how many points and concentration range) was developed, and the resultant detection limit was 0.2 ng/mL. The prepared samples were run in LC-MS and analyzed against the standard curve. For sample preparation, 300 µL of plasma were thawed and spiked with 10 µL of naltrexone (100 µg/mL) as an internal standard. Then, 900 µL of cold acetonitrile was added to precipitate plasma proteins. The samples were vortex-mixed and centrifuged for 10 minutes at 15,000 rpm at 4°C. Upon centrifugation, the supernatant was carefully transferred to the glass tubes and completely evaporated with nitrogen gas at 45°C until all the ACN was evaporated. Dry residues were reconstituted in 120 µL of a mixture containing methanol/water (95:5, v/v), vortex mixed for 5 min, and then filtered using a 0.22 µm hydrophobic filter. The samples were then injected into the LC-MS with Phenomenex LC column (Kinetex 1.7 µm PFP 100 Å, with dimensions of 100 mm x 2.1 mm); the mobile phase consisted of a gradient ranging from 55% to 90% LC-MS acetonitrile (0.1% TFA) and LC-MS water (0.1% TFA). The separation was achieved at a 0.4 mL/min flow rate over 8 minutes. A non-compartmental analysis (NCA) was performed to calculate pharmacokinetic parameters based on the plasma concentration-time profiles. The absolute bioavailability was determined using the following equation:

$$\text{Bioavailability (F\%)} = \frac{AUC_{tested\ route} \times Dose_{IV}}{AUC_{IV} \times Dose_{tested\ route}} \quad (1)$$

#### ***1.2. Fentanyl self-administration training***

After undergoing a five-day recovery period post-surgery, rats were trained to tap on the right lever to receive a 3.2 µg/kg/infusion of fentanyl. The training program utilized an initial fixed-ratio 1 (FR1) schedule with a 20-second timeout between infusions, and the sessions were for 2-

h daily as previously outlined [1]. The fixed ratio represents how many times a rat must press the lever to receive a fentanyl infusion. At the beginning of each session, the rats received a non-contingent fentanyl infusion, followed by a 60-second timeout period. The extension of the right lever and the activation of a green LED light above the lever indicated the availability of fentanyl. After successfully meeting the response requirements, the lever was retracted, and the green light was turned off. Once the rats had earned more than ten fentanyl infusions for a minimum of three days, the schedule was gradually transitioned from FR1 to FR2, FR3, and eventually to FR5. The rats remained on the FR5 schedule for at least five days before being trained to respond to the other lever for liquid food.

#### *1.3. Food-maintained responding training*

Rats were trained to press the left lever for a 5-second presentation of liquid food. The training followed an initial FR1/20-second timeout schedule of reinforcement during daily 2-hour behavioral sessions, as previously described [2,3]. A red LED light above the lever indicated the availability of liquid food. After earning more than 100 food reinforcers, the schedule was raised to the FR5 for at least two days.

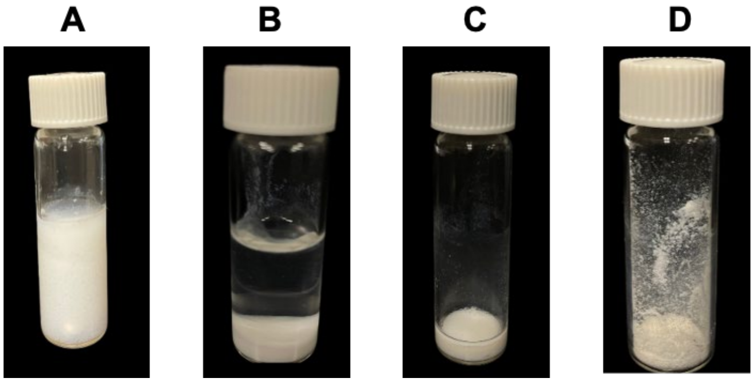


**Fig. S1.** Visual representation of HIP formation. **(A)** nor-LAAM and pamoic acid precipitate complex formation. **(B)** After complex centrifugation. **(C)** Removal of the supernatant in the complex. **(D)** Freeze and dried complex.

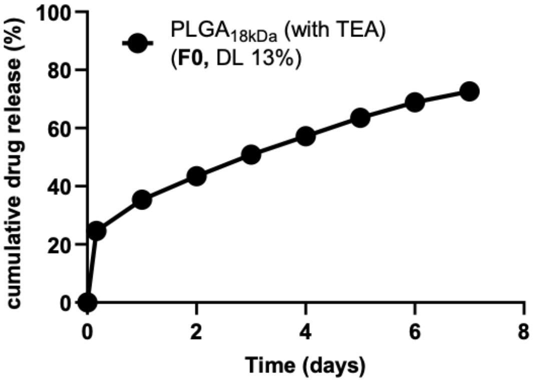


**Fig. S2.** *In vitro* drug release of nor-LAAM-MP prepared using Triethylamine (TEA) without HIP. Mean±SEM (n = 3 repeats).

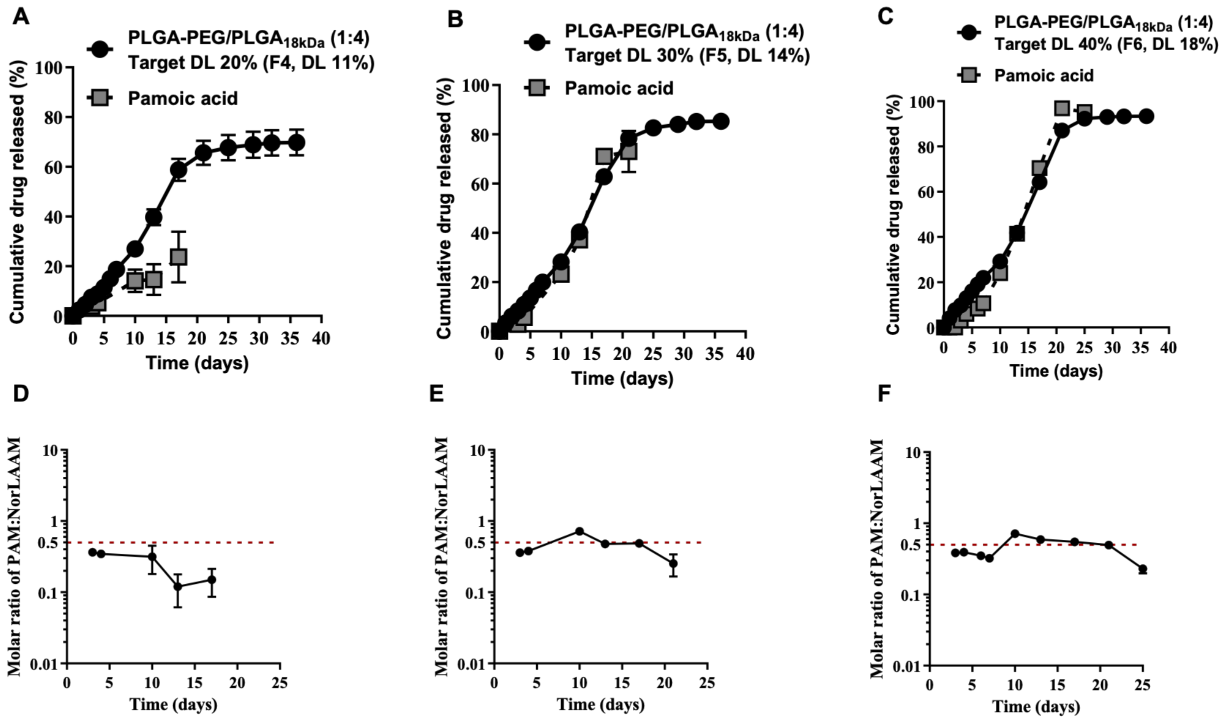


**Fig. S3. (A-C)** In *vitro* release profile of nor-LAAM-MP (F4, F5, and F6) showing the release of nor-LAAM and pamoic acid. **(D-F)** showing the molar ratio of nor-LAAM and pamoic acid. Mean±SEM (n = 3 repeats).

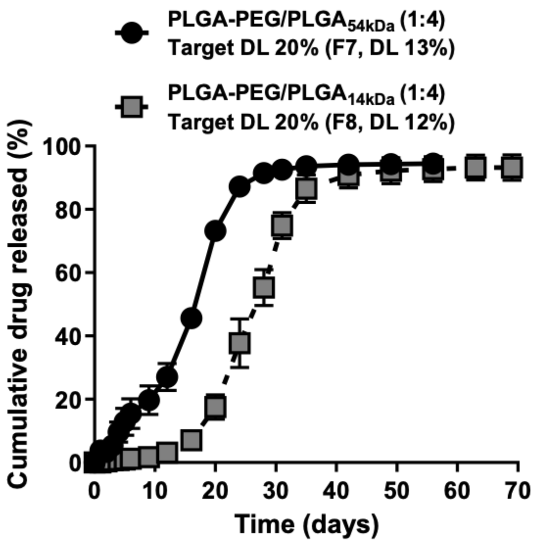


**Fig. S4.** *In vitro* drug release of nor-LAAM-MP-F7 and F8. Mean±SEM (n=3 repeats).

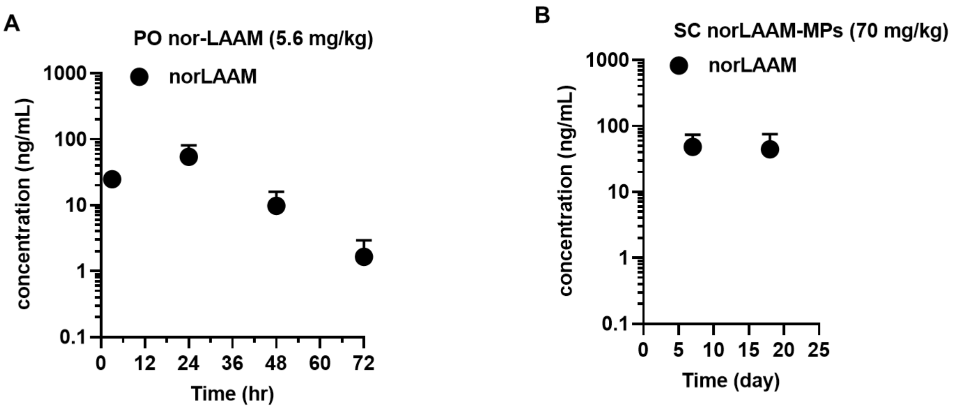


**Fig. S5.** *In vivo* pharmacokinetics profile in rats after administering oral nor-LAAM (n=3) and nor-LAAM microparticles (MPs) (n=2).

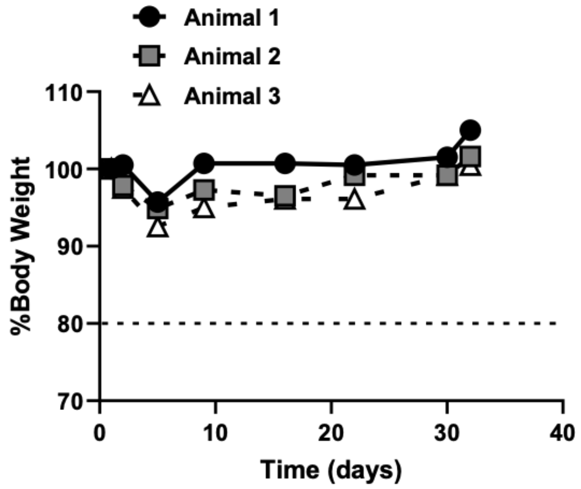


**Fig. S6.** In vivo pharmacokinetics in rabbits demonstrated that nor-LAAM-MP-F4 did not cause a decrease in body weight below 80%.

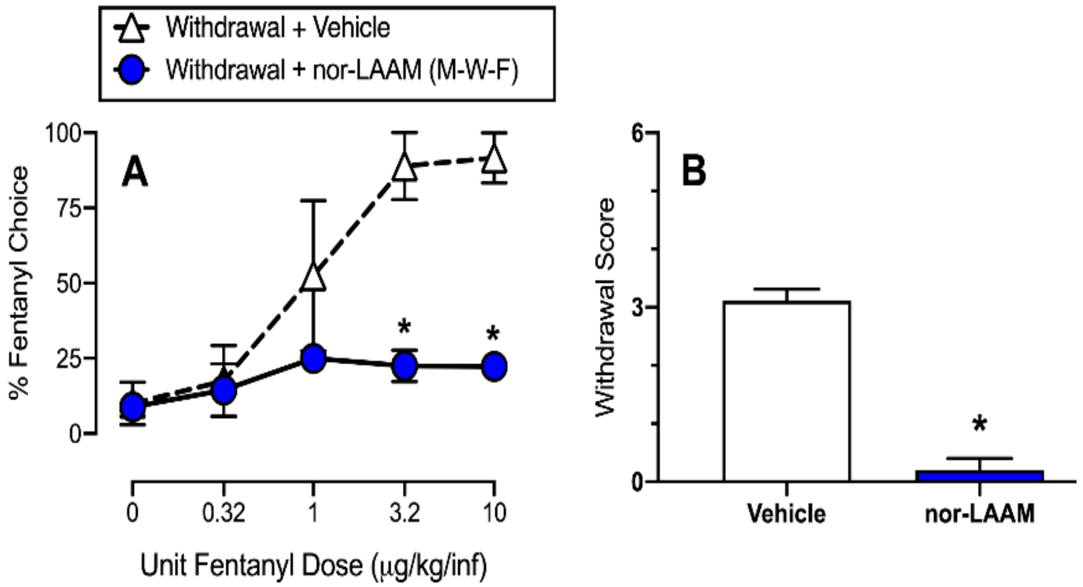


**Fig. S7.** Repeated treatment with norLAAM oral solution (5.6 mg/kg) 3 times a week on xxx rats. (**A)** Oral norLAAM solution decreased fentanyl choice and (**B**) opioid withdrawal somatic signs in opioid-dependent rats. Data are represented as mean±SEM (n=3). * P<0.05.

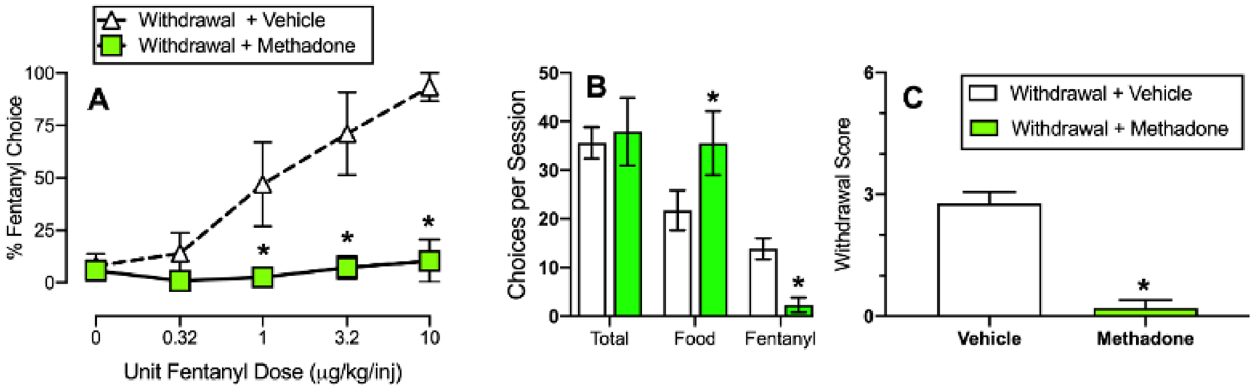


**Fig. S8.** Daily administration of methadone (5.6 mg/kg, 3 times per day). (**A**) Oral methadone solution decreased fentanyl choice. (**B)** shows that chronic methadone significantly increased food choiceand significantly decreased fentanyl choices ($F_{2.2,21.5}$=16.1, p<0.0001). **(C)**: chronic methadone treatment significantly decreased somatic opioid withdrawal signs compared to vehicle (Mann-Whitney test, p<0.0001). (*p<0.05)

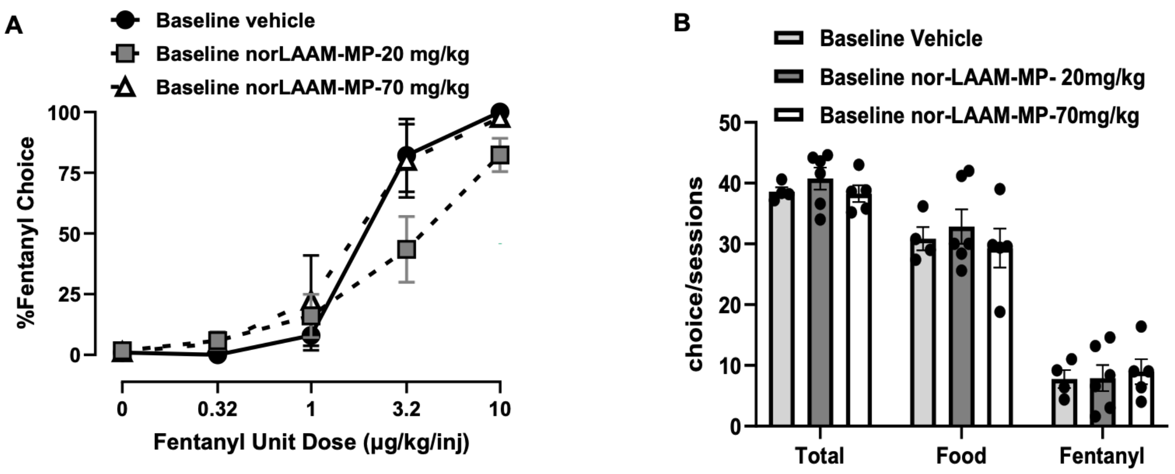


**Fig. S9.** Baseline on fentanyl-vs-food choice before extended fentanyl self-administration. (**A**) percent fentanyl choice in the vehicle (n=4), norLAAM low dose (20 mg/kg) (n=6), and high dose (70 mg/kg) (n=5) treated groups. (**B**) Number of choices completed per session. Data are represented as mean±SEM.

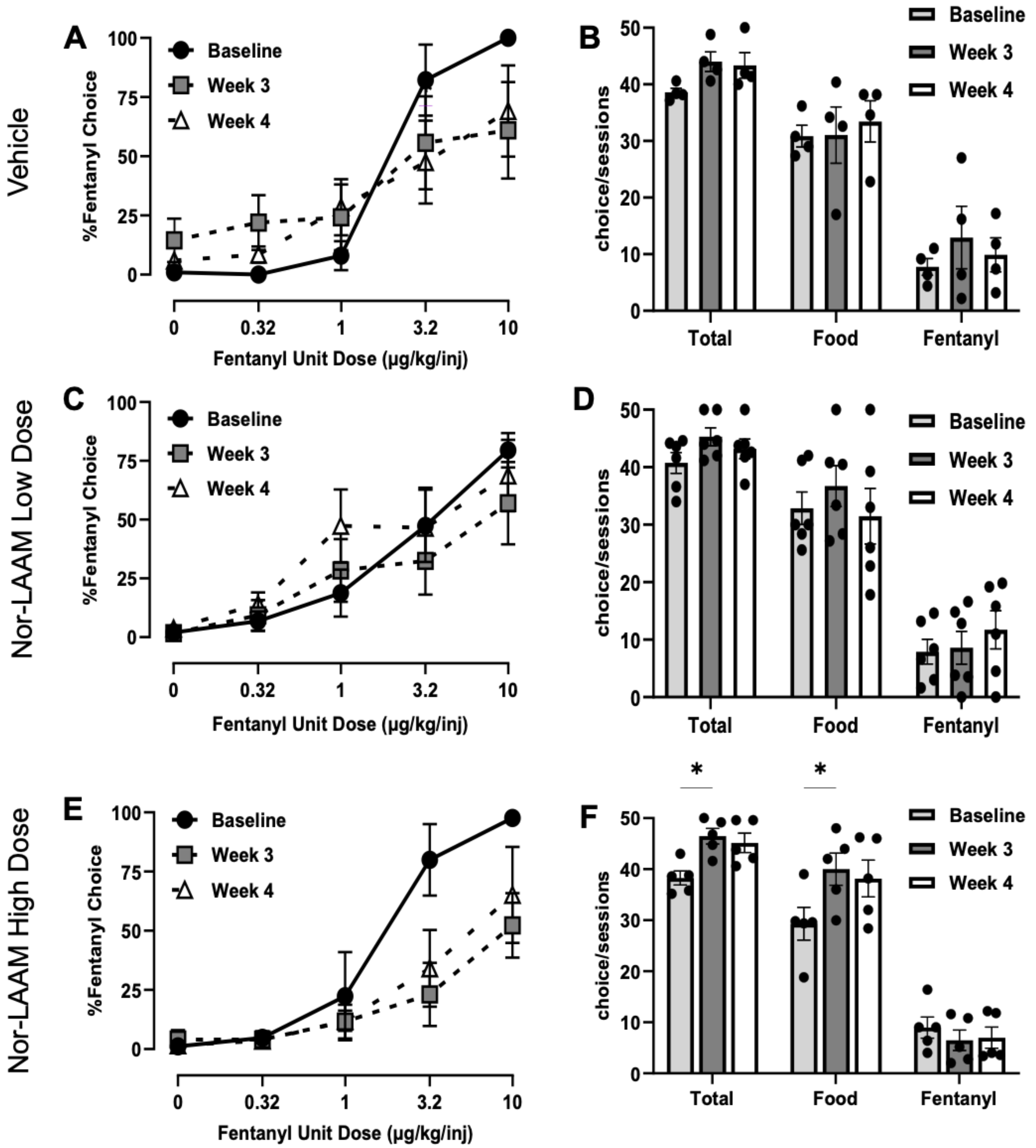


**Fig. S10.** Effects of a single 1-month subcutaneous injection of vehicle (n=4), low-dose norLAAM (20 mg/kg) (n=6), and high-dose norLAAM (70 mg/kg) (n=5) on fentanyl-vs.-food choice in opioid-dependent rats. (**A, C, E**) effect on percent fentanyl choice in weeks 1 and 2. (**B, D, F**) Number of choices completed per session in weeks 1 and 2. Data are represented as mean ± SEM. *symbolize significant difference relative to baseline $p < 0.05$. # symbolizes a significant fentanyl unit dose and time interaction.

[3] Opioid Withdrawal Produces Sex-Specific Effects on Fentanyl Versus Food Choice and Mesolimbic Transcription Supplement 1, n.d.
